# Throwing dice, but not always: bet-hedging strategy of parthenogenesis

**DOI:** 10.1101/2025.11.05.686691

**Authors:** Yanchao Chai

**Affiliations:** Marine Science and Engineering College, Nanjing Normal University, 1 Wenyuan Road, Nanjing 210023, China; Hainan University, 58 People Road, Haikou 570228, China

**Keywords:** Parthenogenesis, Rotifer, Family pedigree, Life history strategy, Evolutionary fitness

## Abstract

Parthenogenesis, clonal propagation by only female, is a common asexual reproduction model. Without sexual gene recombination, it is hypothesized that the deficiency of genotypic variation and accumulation of deleterious mutations reduce the fitness of parthenogenetic lineages confronted with environmental fluctuations, which is also regarded as evolutionary dead end. There should be specific life-history strategies to explain why parthenogenesis has been existing successfully. We constructed a family pedigree for rotifer spanning six generations, comprising 1200 individuals with identical genetic background in uniform condition, tracing back to the inception of parthenogenesis from single dormant egg. The individual fitness represented by lifespan and fecundity exhibits rich variation, and seems to be determined before birth by maternal stochastic investment among clutches regardless of maternal aging. Alike to “Do not put all your eggs in one basket”, this bet-hedging strategy spreads risk of environmental unpredictability. Despite the absence of sexual recombination, the phenotypic fitness failed to achieve fixation and heritability, instead demonstrating transgenerational compensation and trade-offs phenomenon. More siblings mean less children, and vice versa. This can be regarded as intrinsic and innate non-density-dependent self-regulation strategy of population, as limited by the conservation of disposable energy for allocation among offspring. Those strategies are conducive to explain the adaptability of parthenogenesis in evolution.

## Introduction

The fitness for population maintenance is the ultimate aim in evolution, illustrated by successive generation-passage through reproduction^1, 2^. The co-existence of different reproductive models (e.g., asexual vs. sexual) brings controversy around which one is “jack-of-all-trade” confronting with environmental fluctuations^3-5^. The variable genotype then phenotype become countermeasure for this selection pressure. Due to the deficient gene recombination and accumulation of deleterious mutations, the asexual reproduction is always regarded as an evolutionary “dead end”^3^. Paradoxically, asexual lineages are as common as sexual in many ancient organism lineages such as rotifers^6^, which requires more explanations on the adaptive reproductive strategies.

Phenotypic plasticity and bet hedging have been regarded as main strategy for coping with predictable and unpredictable environmental changes, respectively^7^. For asexual lineages, non-genetic polymorphism, a typical plasticity, needs to be triggered by special environmental signal^7, 8^. Compared to predictable varieties (i.e., periodic or recurring changes), the unpredictable one makes more survival risks and selection pressures. Bet hedging occurs when a single genotype produces fitness variance in offspring in advance of future unpredictable conditions^9, 10^. Alike to “Do not put all your eggs in one basket”, bet hedging is bound to the randomness on maternal resource allocation to max long-term fitness of population^11^. The resource allocation can be directly reflected in fecundity and longevity of offspring, namely individual fitness^12^. It has been shown that the joint strategy mixing plasticity and bet hedging can be more successful, and randomly decides which one to adopt by “coin-flipping” way^11^. But, to some extents, this strategy depends on predictable cues. For asexual lineages, the direct empirical evidence on pure bet-hedging potential in advance of environmental unpredictability is still scarce. On the contrary, the effect of maternal age, a regularity in reproduction, seems to limit the randomness of fitness allocation^13^.

In rotifers, asexual and sexual lineages coexist facultatively or obligately, which provides ideal materials for their evolution^7^. In parthenogenetic population, sexual mixis occur periodically to produce resting egg diapause against harsh conditions, but reduce population size, which spreads long-term risks at the expense of current benefits^8^. The variance on diapause-related traits has been regarded as the bet hedging of parthenogenesis in confront environmental fluctuation^8^. Nevertheless, this strategy still depends on sexual reproductive, which is impractical in some obligate asexual lineages. Other types of bet hedging strategies based on randomness may exist to support fitness of parthenogenesis. Therefore, we established parthenogenetically a clonal family (6 generations and 1200 individuals) of *Brachionus calyciflorus* cultivated individually in uniform condition (see Methods for details), and tracked individual longevity and fecundity. The potential reproductive strategies could be explored based on random variability within single genotype.

## Materials and methods

The cloned strain of rotifer *Brachionus calyciflorus* originated from a resting egg, onset of facultative parthenogenesis. After hatched in 24-well plate with culture medium (96 mg NaHCO_3_, 60 mg CaSO_4_, 60 mg MgSO_4_ and 4 mg KCl in 1 L distilled water) including 5×10^6^ cell/ml *Chlorella pyrenoidesa*, the stem individual (F0) was observed every 12 hours, and every newborn larva (F1) was taken out and transferred to single well with 1 mL new culture medium to avoid peer pressure. Repeated this process until population size reached 1200 individuals span 6 generations (Fig. 1). The culture condition was kept constantly in illumination incubator with 16L:8D light schedule with light intensity of 4000 Lux at 25°C. The individual offspring number, birth and death times were recorded.

**Fig. 1.**
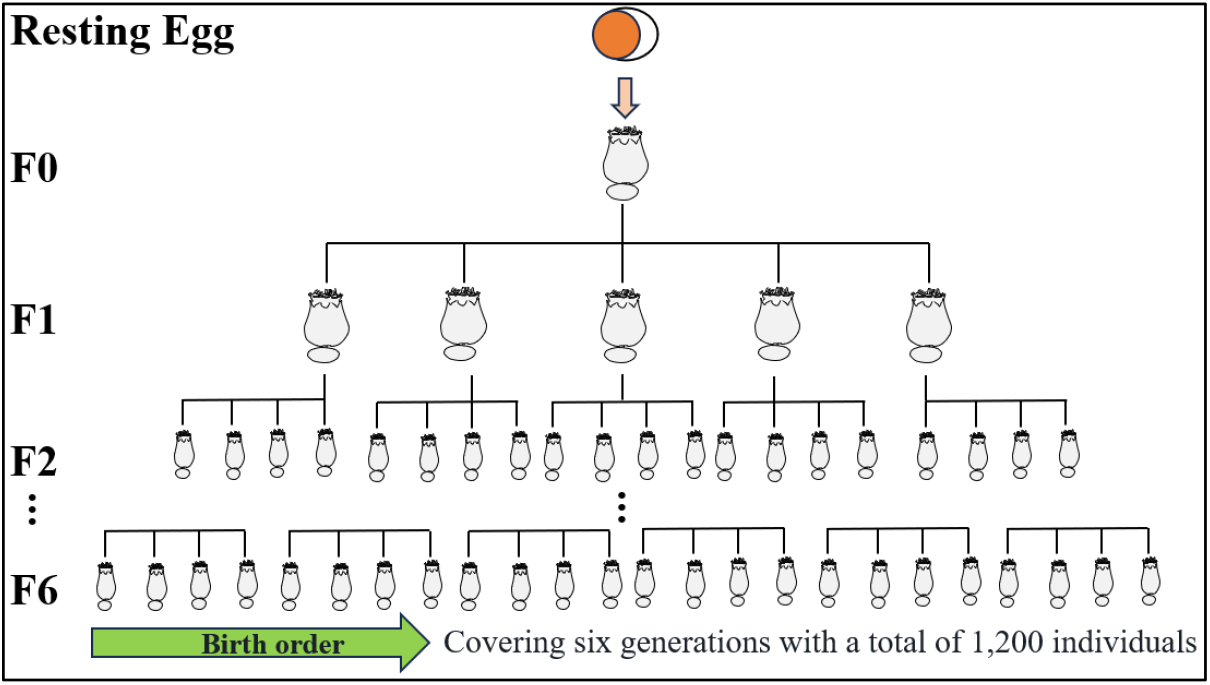
Establishment of parthenogenetic family pedigree

## Results and discussion

Although there was identical gene background, individuals’ lifespan and fecundity exhibited high variability with 39% and 65% coefficient of variation under same and stable environment without competitive pressure (Fig. 2). This reflects evident and huge potential for phenotypic plasticity in the parthenogenetic cloned population.

**Fig. 2.**
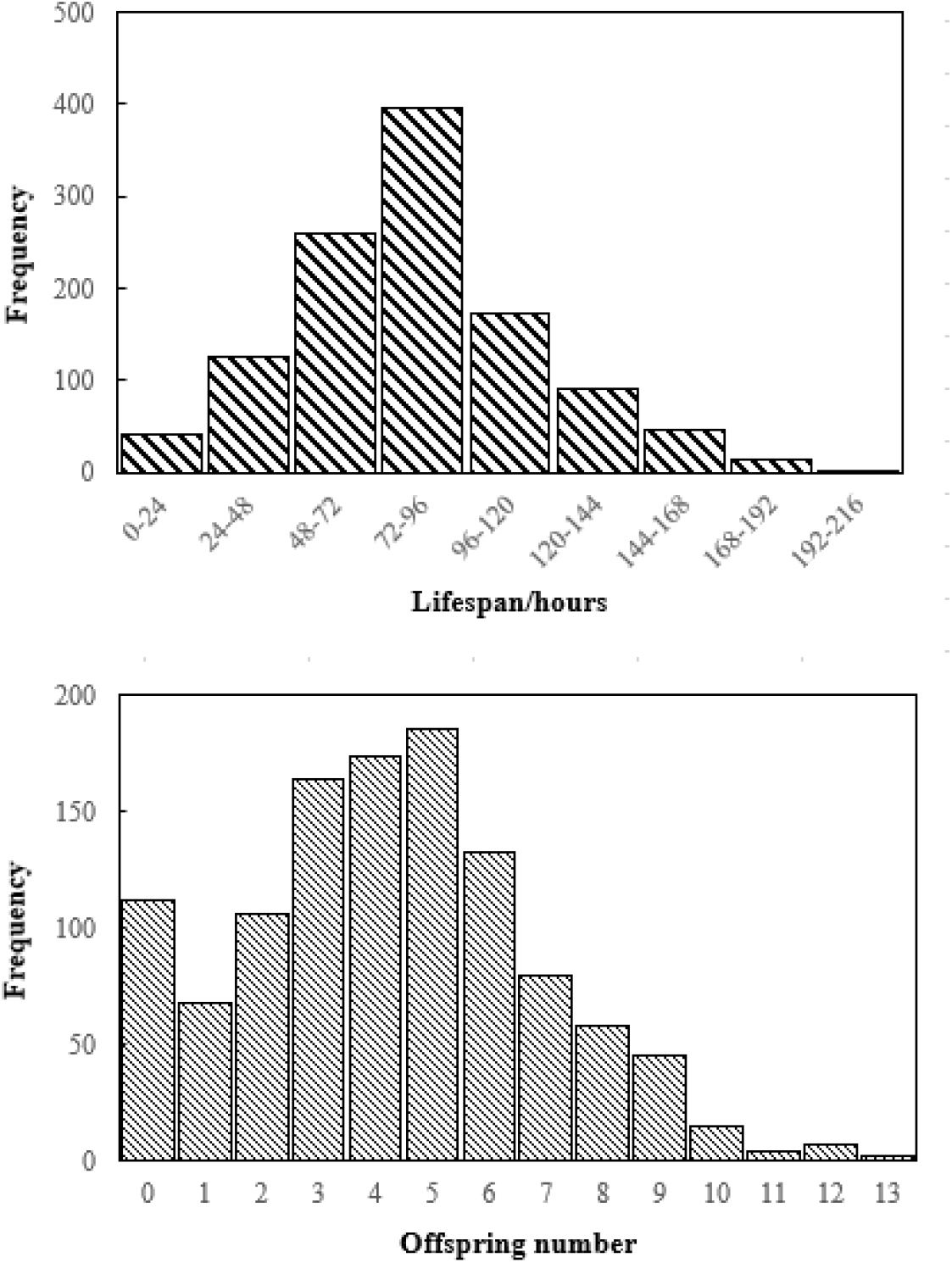
Histogram of individual lifespan and fecundity distribution

**Fig. 2.**
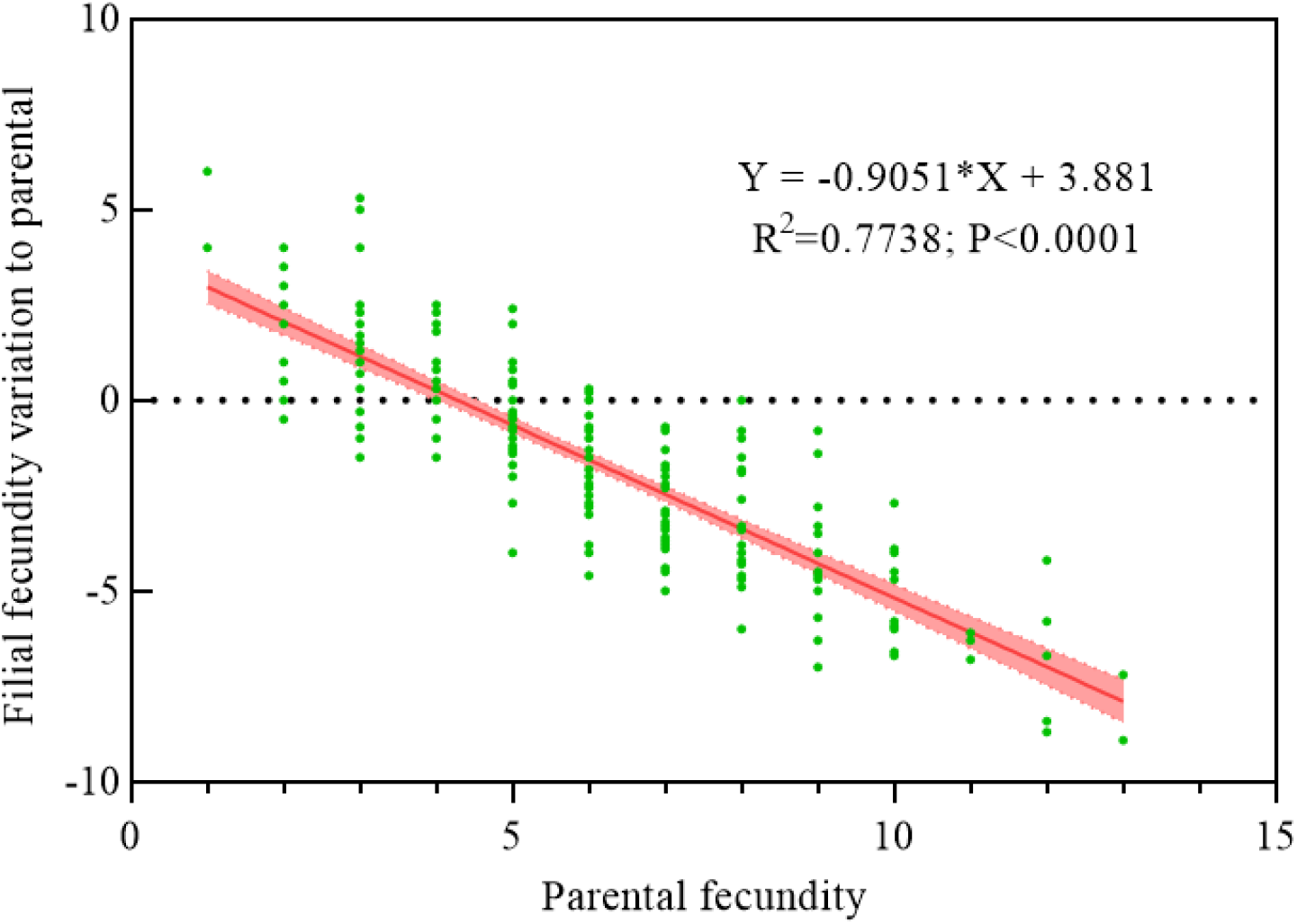
Correlation between parental fecundity and variation of filial fecundity (difference between average fecundity of filial individuals and their parental fecundity)

There was a positive correlation between lifespan and fecundity (Fig. 3). Occasionally, under the premise of conservation of individual total disposable capacity, lifespan and fecundity show negative correlation due to trade-offs of energy allocation among survival, growth and reproduction. That is how organisms contain survival by suppress reproduction confronting with stressed conditions. In consideration of same and comfortable culture conditions in this study, it is logical to deduce that individuals were endowed with different capacity before birth, namely, fitness generally represented by lifespan and fecundity. It is like innate determinism theory based on maternal energy allocation.

**Fig. 3.**
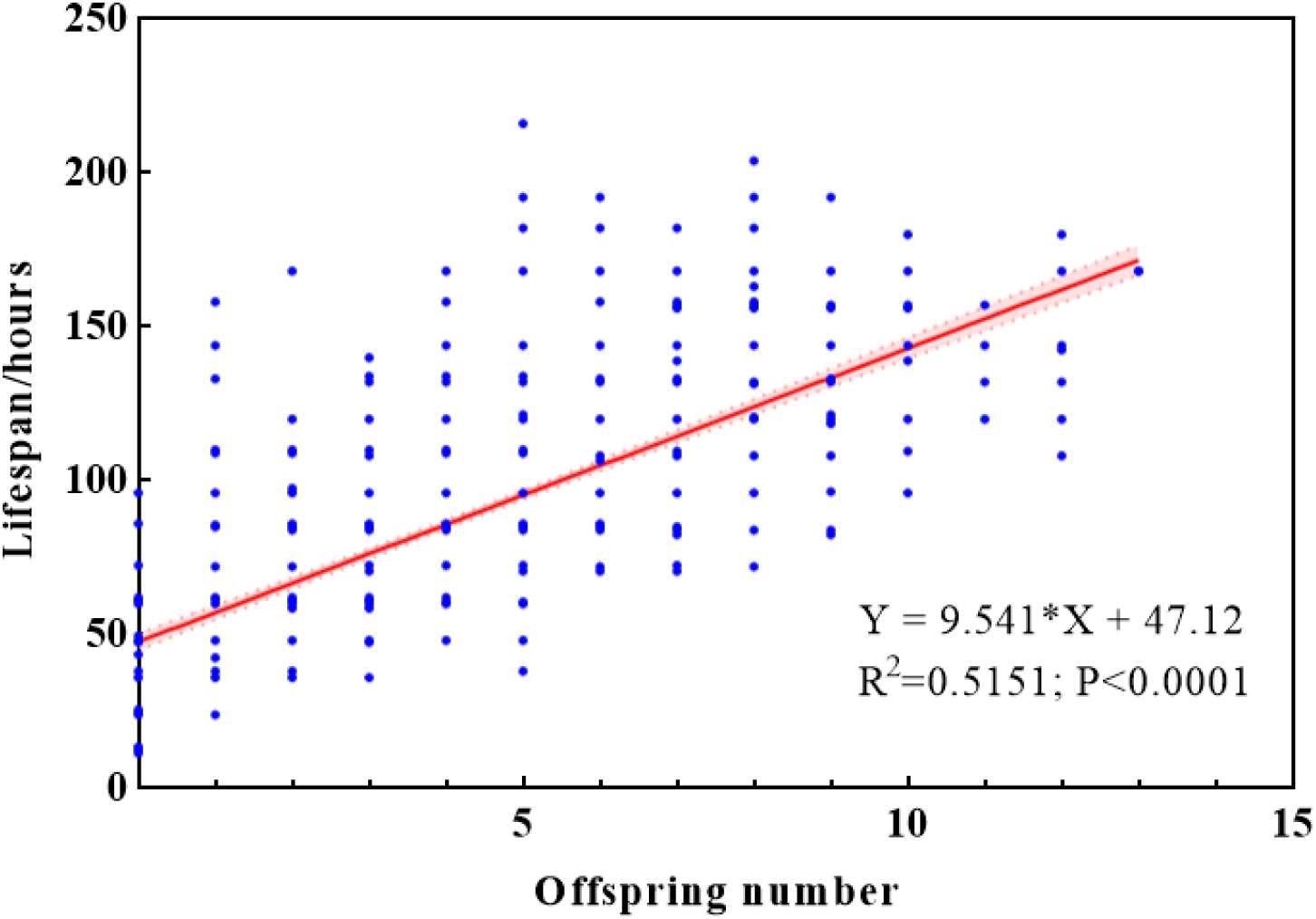
Correlation between lifespan and fecundity

The maternal energy allocation among siblings showed random pattern, as those individuals from different clutches of same mother had similar offspring number distribution regardless of maternal aging effect. And the fecundity variation within clutches could explain variation of entire population (Fig. 4). The maternal random investment to children, like bet-hedging strategy, attributes to the phenotypic plasticity.

**Fig. 4.**
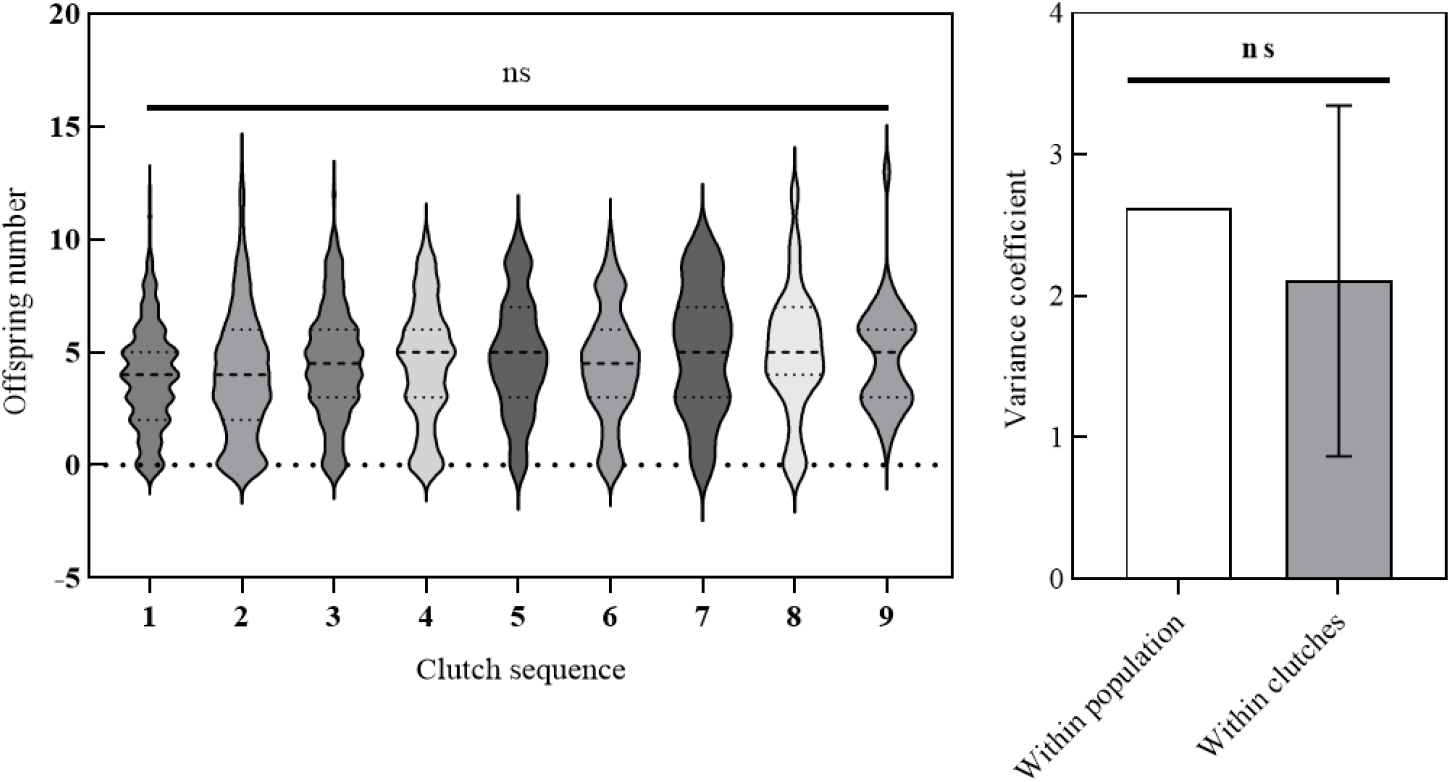
Fecundity variation across clutches

The trait of fecundity characterize by maternal investment could not be inherited by the next generation, as there was no evident correlation between maternal and filial fecundity (Fig. 5). As an advantage of asexual reproduction, it is assumed that the excellent traits can be fixed and inheritable owing to the absence of gene recombination. Nevertheless, this hypothesis seems to be invalid for parthenogenesis, and plasticity still dominates intergenerational transmission of phenotype overriding genotype.

**Fig. 5.**
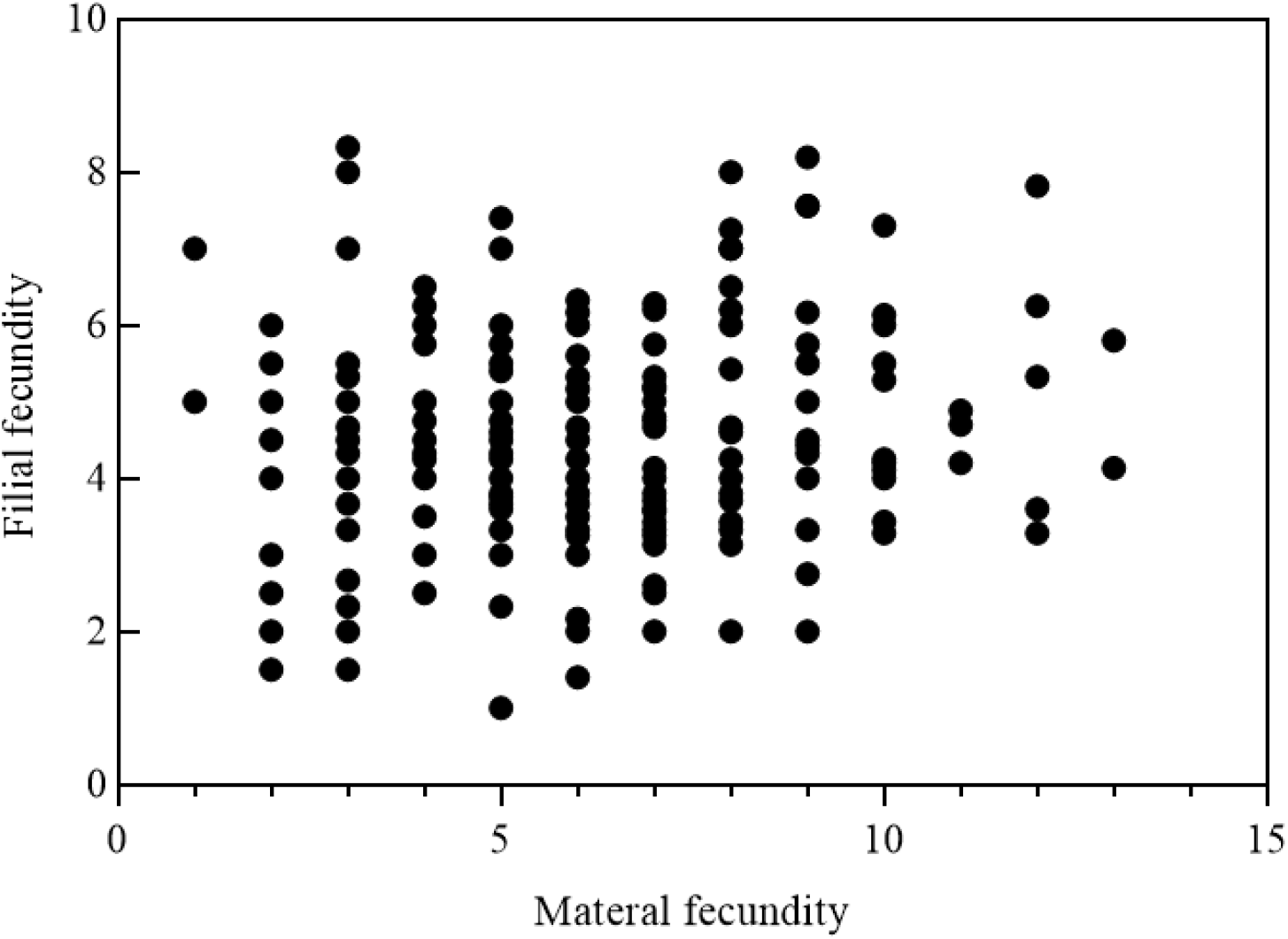
Correlation between parental and filial fecundity

This phenotypic plasticity was not entirely random, but which was control by the transgenerational trade-offs (Fig. 5). To put it another way, if mother produces excessive children, those children’s fecundity will decrease, namely, there is a compensation point. In this study, the point value ranges from 4 to 5, which corresponds with the population fecundity histogram (Fig. 1). Thus, this trade-offs response can be interpreted as a mechanism for population self-regulation coping with unpredictable environmental shocks. The suppressed fecundity under harsh conditions can be compensated in offspring for population restoration once upturn. Meanwhile, excessive reproduction will inhibit fecundity in next generation as a self-thinning to weaken intraclonal competition.

